# Spatial distribution of mammography adherence in a Swiss urban population and its association with socioeconomic status

**DOI:** 10.1101/404673

**Authors:** José Luis Sandoval, Rebecca Himsl, Jean-Marc Theler, Jean-Michel Gaspoz, Stéphane Joost, Idris Guessous

**Author notes:** Correspondence: Prof Idris Guessous and Dr Stéphane Joost, Unit of Population Epidemiology, Department of Community Medicine, Primary Care and Emergency Medicine, Geneva University Hospitals, Rue Gabrielle-Perret-Gentil 4, 1205 Geneva, Switzerland, and. These authors contributed equally (José Luis Sandoval, Rebecca Himsl).

## Abstract

**Purpose:** Local physical and social environment has a defining influence on individual behaviour and health-related outcomes. However, it remains undetermined if its impact is independent of individual socioeconomic status. In this study, we evaluated the spatial distribution of mammography adherence in the state of Geneva (Switzerland) using individual-level data and assessed its independence from socioeconomic status (SES).

**Methods:** Geo-referenced individual-level data from the population-based cross-sectional Bus Santé study (n = 5,002) were used to calculate local indicators of spatial association (LISA) and investigate the spatial dependence of mammography adherence. Spatial clusters are reported without adjustment; adjusted for neighbourhood income and individual educational attainment; and demographic factors (age and Swiss nationality). The association between adjusted clusters and the proximity to the nearest screening centre was also evaluated.

**Results:** Mammography adherence was not randomly distributed throughout Geneva with clusters geographically coinciding with known SES distributions. After adjustment for SES indicators, clusters were reduced to 56.2% of their original size (n = 1,033). Adjustment for age and nationality further reduced the number of individuals exhibiting spatially dependent behaviour (36.5% of the initial size). The identified SES-independent hot spots and cold-spots of mammography adherence were not explained by proximity to the nearest screening centre.

**Conclusions:** SES and demographic factors play an important role in shaping the spatial distribution of mammography adherence. However, the spatial clusters persisted after confounder adjustment indicating that additional neighbourhood-level determinants could influence mammography adherence and be the object of targeted public health interventions.

## INTRODUCTION

Breast cancer is the most common neoplasm in high-income countries [1] and mammography screening has been shown to contribute to its early detection. Mammography screening, together with improvements in disease management, has supported a decrease in breast cancer mortality over the last decades. [2-3] Several studies have identified socioeconomic inequalities in mammography screening adherence and noted the potential impact on the development and persistence of socioeconomic inequalities in breast cancer mortality and morbidity. [4-6]

Mammography screening adherence, like other health-related outcomes, can be affected not only by individual factors (e.g. age, income and education) but also by environmental characteristics, such as neighbourhood context. [7] Physical characteristics of the neighbourhood, including infrastructure quality and housing conditions, may influence the health of its inhabitants. [8] Furthermore, individual social capital and social networks can contribute to different patterns of health outcomes. [9-11]

Studies investigating the influence of neighbouring effects on mammography adherence are mainly ecological in nature and have considered artificial geographic groupings (neighbourhoods, counties, zip codes, etc) rather than the geographic distance between individuals.

Through the use of spatial analytic methods that take geographic distance into consideration, clusters of individuals sharing similar health behaviours and characteristics can be identified [12-13] in order to tailor public health interventions to the populations in need. In addition, spatial clustering may uncover links between spatial proximity and health outcomes that are independent of socioeconomic status (SES) and other individual characteristics that would otherwise be missed if the spatial context was not considered.

We have previously identified SES inequalities in mammography screening adherence in the state of Geneva, Switzerland, and have studied their temporal dynamics using data from a yearly cross-sectional study spanning 22 years. [6]

In the present study, we first aimed at determining if geographical clusters of mammography non-adherence exist in the population of Geneva and, second, to determine if these were independent of individual demographic and socioeconomic characteristics.

## METHODS

### Participants

We included data from female participants in the Bus Santé study who were between the ages of 50 and 74 and had no history of breast cancer. Details of the Bus Santé study and its sampling strategy have been described elsewhere [14]. In brief, a representative stratified sample of the Genevan population (Switzerland, ∼500,000 inhabitants) has been collected every year since 1993. Non-institutionalised residents between the ages of 35 and 74 (from 20 to 74 after 2011) were selected from an annual residential list compiled by the local government and were then subjected to stratified random sampling based on gender and 10-year age strata. Data were sourced from self-administered, standardised questionnaires that concentrate on individual sociodemographic characteristics and disease risk factors. Geographic coordinates were derived from participants’ residential addresses. Participation rates ranged from 55 to 65% and were lower in 2005 and 2006 due to a concurrent study that shared resources with the Bus Santé study but did not target the same population.

This study was approved by the Institute of Ethics Committee of the University of Geneva and written consent was obtained from all participants.

Participants were excluded from the analysis if either geographic data (n=164, 3.1%) or individual-level confounder information (n=168, 3.1%) were missing. Missing data were assumed to be missing completely at random. A total of 5,002 participants were included in this study.

### Variables

Geographic coordinates of each participant’s postal address were obtained using the IDPADR, a unique and permanent street-number identifier used by the State of Geneva to manage the addresses of buildings on its territory. The dataset used in this analysis was geocoded by matching the Bus Santé participants’ IDPADRs with those given in the State’s comprehensive spatial database (www.ge.ch/sitg/donnees/demarche-open-data).

A binary outcome variable, mammography non-adherence, identified women that had never had a mammogram. Educational attainment was considered in 3 levels as in Huissman et al. [15]: primary (no primary school certification or professional apprenticeship), secondary (completed secondary education or professional apprenticeship) and tertiary (university degree). Area income level in CHF/year was obtained from the 2013 Geneva Census (www.ge.ch/statistique, Geneva Statistics Office) and used as a proxy for individual income (Figure S1). In May 2018, 1 CHF corresponded to approximately 1 USD, 0.76 GBP and 0.86 EUR. Nationality (Swiss or other) and participant age (continuous) were also considered as confounders. Median revenue was not reported for statistical units with less than 20 inhabitants (n =94). Of the 5,002 participants in this study, 78 (1.6%) resided in these units. In order to maintain the spatial contiguity of the dataset, replacement by mean was used to fill in missing values (mean = 73,191.82 CHF).

Differences in confounding variables between participants who had had a mammography screening and those who had not were estimated. Two-tailed t-tests were used to assess the between-group differences in age and median revenue, ANOVA was used for education level, and a chi-square test was used to evaluate the significance of Swiss nationality.

R Statistical Software (version 3.3.1, R Foundation for Statistical Computing, Vienna, Austria) was used for all statistical procedures. An α value of 0.05 was used for all statistical testing.

### Confounder-adjusted spatial analysis

In order to perform spatial analysis on mammography adherence adjusted for several potential confounders, we fit the data with a logistic multivariate regression model and extracted the Pearson residuals. Geographic clustering was performed on the residuals whereby it is assumed that, after having adjusted for confounding factors, any spatial association exhibited by the residuals can be predominantly attributed to external spatially dependent factors.

Two different models were fit: one to adjust mammography non-adherence for education and income; the second, not only for the SES variables, but also for participant age and nationality.

### Spatial analysis

We used the software GeoDa (1.10.0.8) to calculate the Local Indicators of Spatial Association (LISA) [16] for adjusted and unadjusted mammography adherence. The local indicators constitute a decomposition of the global Moran’s I index [17] into observation-level indices which can measure spatial dependence and evaluate the existence of localized spatial clusters. A local I statistic, Z-score and p-value were computed for each observation by assessing the correlation between the observed outcome (in this case – mammography non-adherence) and the mean behaviour exhibited by neighbouring points within a defined spatial lag. Standardized LISA statistics were then plotted on a scattergram to create five distinct classes: 1) NoAdh-NoAdhh: individuals who have not had a mammogram that live in an area characterized by mammography non-adherence; 2) Adh-Adh: individuals who have had a mammogram and live in an area characterized by mammography adherence; 3) Adh-NoAdh: adherent individuals considered to be outliers residing in a predominantly non-adherence area; 4) NoAdh-Adh: non-adherence individuals considered to be outliers residing in a predominantly adherence area; 5) no spatial dependence. “Adh” and “NoAdh” correspond respectively to the “low” and “high” qualifiers used in Anselin’s original work. [16]

Permutation-based significance testing was used to determine whether an individual’s behaviour will be classified into one of the four spatial clusters or as having no spatial dependence. This was done by means of a large number of random Monte-Carlo permutations, which shift the observed values between the different sample locations. The distribution of the permuted values was compared to the observed LISA statistic, and the significance was calculated as (M + 1) / (P + 1), where P was the number of permutations and, for a positive LISA indicator, M was the number of instances where a statistic computed from the permutations was equal or greater than the observed value; for a negative indicator, M was the number of instances where a permutation statistic was less than or equal to the observed LISA indicator.

The mean distance between each participant and their nearest screening centre is 1,538m and, we chose to use the size of this shared neighbourhood to set the range for the spatial lag used in our spatial analysis. Correspondingly, only results generated using a 1,600m spatial lag are presented here. Nevertheless, the univariate LISA statistic was computed for 13 different spatial lags ranging from 200m to 2,600m in order to assess the influence of neighbourhood size on observed clusters. All lags between 600m and 2,600m showed a similar spatial distribution of adherence and non-adherence clusters.

Maps included in this manuscript report associations at a significance level of p<0.05 given by 999 permutations, where white points identify non-significant LISA statistics.

### Association with proximity to screening centre

We sought to understand if the adjusted mammography adherence patterns could be related to the distance from the nearest mammography centre. This association was evaluated at the global level as well as at the cluster level.

The global level association was assessed by first classifying each participant according to the behaviour exhibited by their respective neighbourhoods; that is each individual is classified as belonging to either an adherence hot-spot or cold-spot. A cold-spot is defined by individuals who were classified as either belonging to the NoAdh-NoAdh or Adh-NoAdh LISA clusters, and a hotspot is composed of individuals who belonged to either the Adh-Adh or NoAdh-Adh LISA clusters. The distance between each individual and the nearest mammography screening centre was then computed, and the mean distance between those belonging to an adherence hotspot or cold-spot and the nearest screening centre was compared using a t-test. The proximity to the nearest screening centre was also considered independently for each spatial cluster. In order to do so, 22 polygons were generated to represent the neighbourhoods characterized by the spatial distribution of mammography adherence. Individuals belonging to each cluster were manually specified so as to satisfy three criteria: 1) individuals within a cluster all resided in either an adherence hotspot or cold-spot; 2) the distance between neighbouring individuals was minimized (i.e. if the difference between two individuals exhibiting the same behaviour was too great, a new cluster would be created); 3) no cluster contained less than three participants. Convex hulls were then used to generate polygons from the categorized individuals. The distance to the nearest screening centre was then calculated from the polygon centroid.

## RESULTS

We included 5,002 participants with a mean age of 60.3±6.8 years - seventy-eight percent (n=3,857, 77.5%) of whom were Swiss. Concerning educational attainment, 28.3% (n=1,417) had primary education while 43.1% (n=2,155) and 28.6 (n=1,430) had secondary and tertiary education, respectively. Mean income was 73,191.82 ± 17,269.28 CHF/year. Twelve percent (n=585, 11.7%) were never screened for breast cancer using mammography. The socioeconomic and demographic characteristics of the participants are summarized according to mammography adherence in Table 1.

**Table 1.**
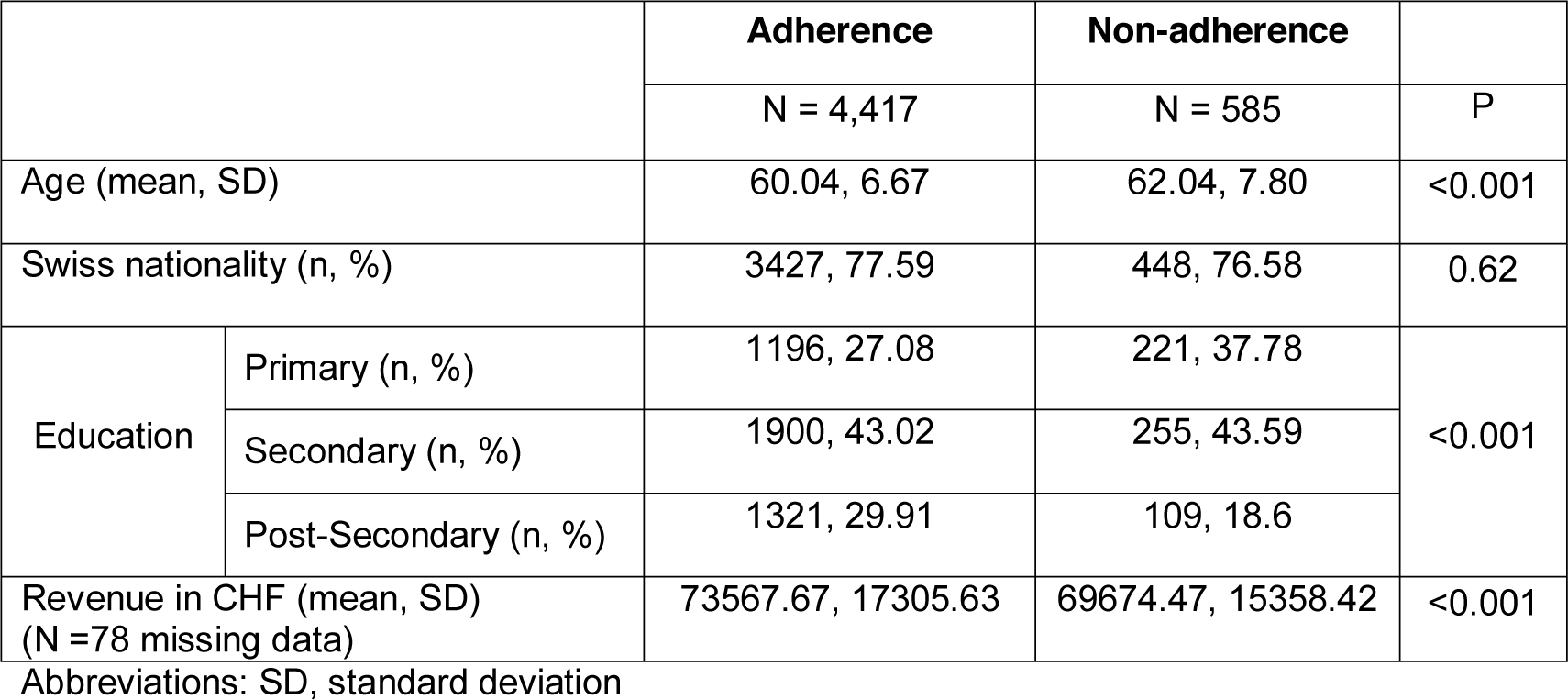
Table 1 summarizes the characteristics of study participants according to whether or not they have undergone a mammography screening.

|  |  | **Adherence** | **Non-adherence** |  |
| --- | --- | --- | --- | --- |
|  |  | N = 4,417 | N = 585 | P |
| Age (mean, SD) |  | 60.04, 6.67 | 62.04, 7.80 | <0.001 |
| Swiss nationality (n, %) |  | 3427, 77.59 | 448, 76.58 | 0.62 |
| Education | Primary (n, %) | 1196, 27.08 | 221, 37.78 | <0.001 |
|  | Secondary (n, %) | 1900, 43.02 | 255, 43.59 |  |
|  | Post-Secondary (n, %) | 1321, 29.91 | 109, 18.6 |  |
| Revenue in CHF (mean, SD)<br>(N =78 missing data) |  | 73567.67, 17305.63 | 69674.47, 15358.42 | <0.001 |

Abbreviations: SD, standard deviation

### Geographical clusters of mammography non-adherence

The unadjusted LISA clusters for the 5,002 participants are shown in Figure 1. Within a 1,600m spatial lag, we observed that the behaviour of the majority of participants did not present spatial dependence (63.3%, n=3,164), 3.6% of participants were classified as NoAdh-NoAdh (n = 178) – that is they had not had a mammogram and resided in an area that showed a higher non-adherence than expected at random. Approximately ten percent of individuals were classified as Adh-Adh (9.7%, n = 485), meaning they had had a mammogram and resided in a neighbourhood that showed a higher adherence than expected at random. Twenty-two percent of individuals belonged to the Adh-NoAdh clusters (22.3%, n = 1,116), and 1.2% to the NoAdh- Adh clusters (n = 58), both of which exhibit discordant behaviours; respectively, these correspond to individuals who had had a mammogram and those who had not but live in areas where their neighbours exhibit the opposite behaviour.

**Figure 1.**
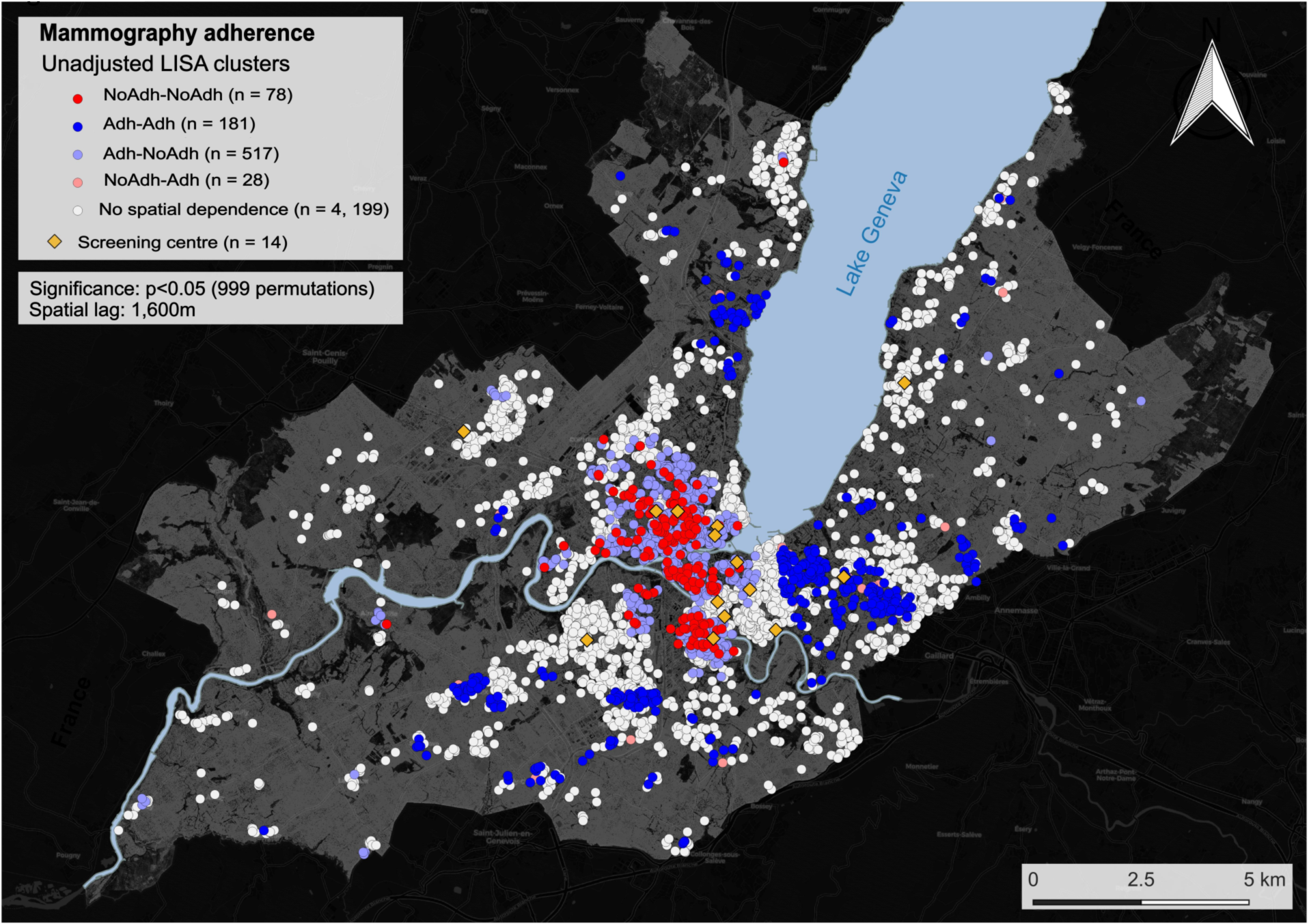
Clusters for mammography non-adherence and adherence in unadjusted spatial analysis. White dots represent sampling locations where no spatial dependence was observed. The following LISA clusters are presented: i) NoAdh-NoAdh: individuals who have not had a mammogram live in an area characterized by mammography non-adherence; ii) Adh-Adh: individuals that had a mammogram and live in an area characterized by mammography adherence; iii) Ad-NoAdh: individual that had a mammogram residing in a predominantly non-adherence area; iv) NoAdh-Adh: individual who have not had a mammogram and considered an outlier residing in a predominantly adherence area. Clustering was performed using a 1,600m spatial lag. The yellow diamonds represent the locations of radiology centres performing mammography screening.

Non-adherence hotspots were preferentially located downtown Geneva and adherence hotspots in the periphery, following the known income distribution for the population of Geneva (Supplementary Figure 1).

Adjusted spatial analysis for known SES indicators (neighbourhood income and education) is presented in Figure 2A. After adjustment for these confounders, spatial independence was observed for 83.9% (n=4,197) of participants’ behaviour, while 1.6% belonged to the NoAdh-NoAdh class (n=78), 3.6% to Adh-Adh (n = 181), 10.3% to Adh-NoAdh (n = 517) and 0.6% to the NoAdh-Adh class (n = 28).

**Figure 2.**
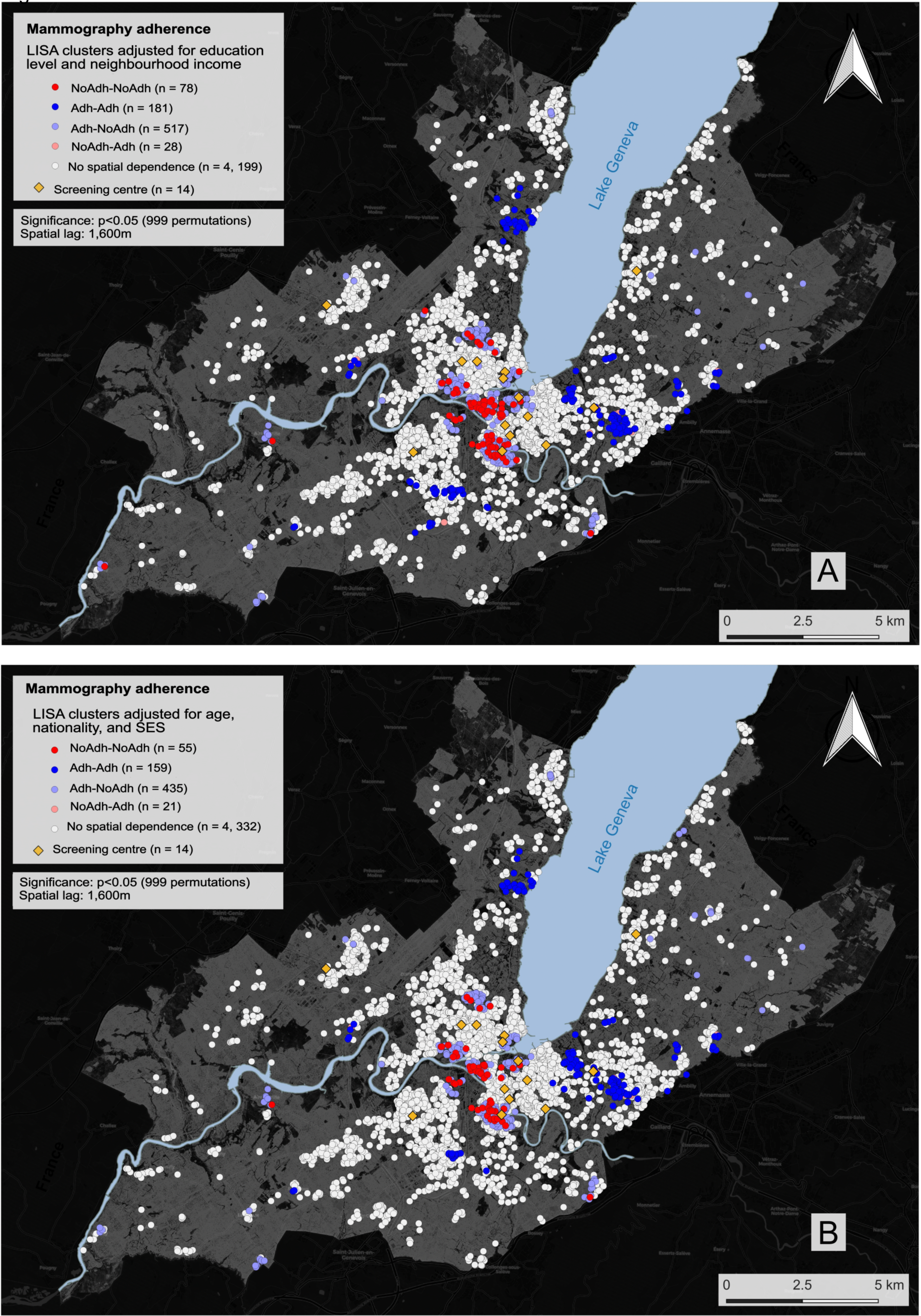
Clusters for mammography non-adherence and adherence (A) adjusted for income and education and (B) income, education, age and Swiss nationality. White dots represent sampling locations where no spatial dependence was observed. The following LISA clusters are presented: i) NoAdh-NoAdh: individuals who have not had a mammogram live in an area characterized by mammography non-adherence; ii) Adh-Adh: individuals that had a mammogram, and live in an area characterized by mammography adherence; iii) Adh-NoAdh: individual that had a mammogram residing in a predominantly non-adherence area; iv) NoAdh-Adh: individual who have not had a mammogram and considered an outlier residing in a predominantly adherence area. Clustering was performed using a 1,600m spatial lag. The yellow diamonds represent the locations of radiology centres performing mammography screening.

Additional adjustment was performed for other confounding variables (age and Swiss nationality) in addition to neighbourhood income and education (Figure 2B). In this model, spatial clustering was further reduced with 86.6% (n = 4,331) of participants’ behaviour not exhibiting spatial dependence. NoAdh-NoAdh clusters were reduced to 1.1% (n = 55); Adh-Adh to 3.2% (n = 159); Adh-NoAdh (n = 435), and NoAdh-Adh to 0.4% (n=21).

In both adjusted analyses, the geographic distribution of non-adherence and adherence hot-spots was analogous to that of the unadjusted analysis, with non-adherence being marked in central Geneva and adherence predominantly in peripheral regions.

Of the 14 screening centres, only 2 were located closer to adherence hotspots than non-adherence, and individuals located in the NoAdh-NoAdh or Adh-NoAdh clusters (or adherence cold-spots) were living on average significantly closer to screening centres than those classified as Adh-Adh or NoAdh-Adh (p<0.001).

## DISCUSSION

Our results show that geographical clusters of mammography adherence can be identified in an urban setting like Geneva, Switzerland. While similar clusters have been previously reported in other circumstances using ecological data [18-19,7,20], we report it for the first time using data at the individual level by considering geographic space as a continuum rather than according to predefined administrative units.

Previous ecological studies, such as Lemke et al., have suggested that clustering was due to low-income and immigrant populations inhabiting neighbourhoods with low mammography adherence [19]. Ecological studies are prone to the ecological fallacy with conclusions about individuals being drawn from group-level analyses; they are based on the mean behaviour of each group and are thereby unable to assess within-group variation. In the epidemiological context, ecological studies tend to consider predefined, administrative, geographic units (neighbourhood, counties, postal code, etc.) which may not reflect the true geographic distribution and heterogeneity of health outcomes and individual characteristics expected in urban settings. Here, clusters based on geographic distance - derived from precisely georeferenced individuals (coordinates of the building’s entrance) rather than administrative divisions - have enabled us to identify regions of mammography adherence and non-adherence, which are determined by individual behaviour and characteristics.

As in other studies, the unadjusted adherence and non-adherence clusters corresponded to areas that are known to be inhabited by people with contrasting SES, suggesting that geographic disparities in SES could explain the observed spatial clustering. While it is known that SES inequalities exist in our population, adjustment for two main SES indicators (education and income) revealed clusters of adherence and non-adherence that were independent of socioeconomic status. It is worth nothing that the clusters that persisted after adjustment for education and median neighbourhood income were significantly smaller; incorporating demographic factors into the adjustment further reduced cluster size to only 36% of the unadjusted clusters indicating that demographic and socioeconomic factors play an important role in determining mammography adherence in the Genevan population. Further, proximity to mammography screening centres seems to be unrelated to the observed spatial clusters.

The existence of SES-independent geographic clusters stresses the need to consider the geographic properties associated with spatial phenomena when studying health outcomes inequalities and devising interventions to address them. This phenomenon may be explained by spillover effects, with spatial externalities occurring when individual knowledge and preferences are transmitted through informal social networks thereby influencing others’ behaviours. [10-11]

In addition, social cognitive theory postulates that behaviour may be influenced by observing the actions of others and their consequences, with each individual being simultaneously a responder and a social stimulus of behaviour, potentially determining the observed SES-independent mammography adherence clusters. [21] This could be the case for mammography screening for which concerns about its efficiency and secondary effects might disseminate through social networks. [22] However, in a study using data from the Framingham Heart Study, Keating et al. found that social networks had a minimal effect on mammography screening behaviour. Social network analysis took into account the influence of siblings, friends and co-workers but the influence of the participants’ geographic context was not explored. [23] While area-level analyses may help identify predefined regions that could benefit from spatially focused interventions, geographical clusters, such as those based on individual data, may be missed if these are scattered across several existing administrative regions.

Our study highlights the importance of considering the geographical distribution of health outcomes at the individual level in order to better tailor public health interventions and proceed to an era of precision public health delivery.[24-25]

### Strengths

Unlike ecological studies, the geographical clustering we propose was defined on the basis of individual data, limiting the potential bias due to ecologic fallacy. This study is, as far as we know, one of the first to use individual geographic coordinates to cluster behaviours related to health outcomes. Furthermore, the availability of sociodemographic data coupled with individual-level geographic data allowed us to exclude that the observed geographic distribution was, entirely, a reflection of differential distributions of SES or other available individual characteristics.

### Limitations

First, similarly to other studies, the adjusted analyses in our study are limited to the confounders available in the Bus Santé dataset, and potential residual confounding cannot be excluded. Second, all data were recorded at the individual level except for income, and to improve the sample size for the SES adjusted spatial analyses we used area-level income as a surrogate for individual income. However, while we identified outcome clusters that corresponded to areas known to show different average income levels, these clusters persisted after adjustment for income, and thus could not be entirely by socioeconomic disparities. Third, mammography adherence was defined as *having ever had a mammography screening* rather than *compliance with local guidelines* (every two years). [26] Data on mammography screening frequency was not available for the whole studied period (1992-2014) and might have overestimated adherence.

## CONCLUSIONS

Using individual-based geographic and sociodemographic data, we have identified SES-independent clusters of mammography adherence and non-adherence in an urban area of Switzerland. Studies focusing on outcome-driven geographical clustering may contribute to better informing decision-makers where to deploy public health interventions in order to help fulfil the promises of precision public health delivery.

## Acknowledgments

The authors are extremely grateful to the Bus Santé study participants who kindly agreed to participate in this study.

## Ethics approval

Institute of Ethics Committee of the University of Geneva

## Author contributions

José Sandoval: Conceptualization, methodology, analysis, interpretation of results, writing and revision of the manuscript. Rebecca Himsl: Conceptualization, methodology, analysis, interpretation of results, writing and revision of the manuscript. Jean-Marc Theler: data collection, interpretation of results, reviewing and editing the final manuscript. Jean-Michel Gaspoz: data collection, interpretation of results, reviewing and editing the final manuscript. Stéphane Joost: Conceptualization, data collection, methodology, analysis, interpretation of results, writing, revision of the manuscript. Idris Guessous: Conceptualization, data collection, methodology, interpretation of results, writing, and revision of the manuscript.

---

Supplementary figure 1 - Distribution of mean yearly income by neighbourhood in the state of Geneva, in CHF. Data were obtained from the 2013 Geneva census (www.ge.ch/statistique, Geneva Statistics Office)

Abbreviations: SD, standard deviation

